# *RAS* Internal Tandem Duplication Disrupts GAP-binding to Activate Oncogenic Signaling

**DOI:** 10.1101/737098

**Authors:** Andrew C. Nelson, Thomas Turbyville, Srisathiyanarayanan Dharmaiah, Megan Rigby, Rendong Yang, John Columbus, Robert Stephens, Drew Sciacca, Getiria Onsongo, Anne Sarver, Subbaya Subramanian, Dwight V. Nissley, Dhirendra K. Simanshu, Emil Lou

## Abstract

Molecular testing of oncogenic *RAS* mutations in colorectal cancer (CRC) have led to increased identification of mutations in patients with CRC. *NRAS*-mutated CRC has not been well characterized because it is less common (<6%) than *KRAS* mutations. Here, we report a novel 10 amino acid internal tandem duplication (ITD) in *NRAS*, which disrupts the switch II domain, in a patient with widely disseminated CRC. Hotspot next generation sequencing of a brain metastasis identified the *NRAS* ITD and a *TP53* missense mutation (p.P275F). Whole exome sequencing of the primary tumor and two metastatic lesions (lung and brain) confirmed that the *NRAS* ITD and *TP53* mutation were conserved between the primary tumor and both metastatic tumors, and identified an additional pathogenic mutation in *CSMD1* (a tumor suppressor gene). Structural biology and biochemical analyses demonstrated that the *NRAS* ITD prevented binding to GAP protein, leading to sustained RAS activation, increased interaction with RAF, and downstream MAPK activation. Additionally, we provide the first crystal structure of the *RAS* ITD. In conclusion, these studies indicate that the *NRAS* ITD was the probable primary driver mutation of this aggressive CRC. Identical or biologically similar ITDs in *NRAS* and *KRAS* may be rare drivers of CRC and other aggressive malignancies.

## Introduction

Colorectal carcinoma (CRC) is a major cause of cancer morbidity and mortality with over 1.8 million new cases diagnosed annually worldwide (1). Nearly 20% of CRC cases are metastatic and incurable at the time of diagnosis (2). Well-characterized driver mutations in *KRAS*, primarily at codons 12, 13, 61, and 146, are identified in 35-40% of CRC cases. An additional 5-10% of CRC cases harbor a mutation in *NRAS* at functionally identical codons. *RAS* mutations are associated with decreased response to EGFR inhibition and increased incidence of distant metastasis (3). Limited *KRAS* hotspot mutation testing in CRC has been the standard-of-care as a predictive biomarker to select *KRAS*-wild type tumors that will likely respond to anti-EGFR monoclonal antibody therapy (4).

Recent advances in clinical genomic profiling have expanded testing for a broader spectrum of mutations in *KRAS, NRAS, and BRAF* in patients with metastatic CRC. This expansion has led to the increased identification of rare or previously uncharacterized *RAS* mutations. In particular, the clinical characteristics of *NRAS*-mutated CRCs have not been well established because of their relatively low prevalence. Furthermore, the biologic properties of *NRAS*-mutated cancers do not necessarily overlap with the properties of *KRAS*-mutated cancers (5–7). Thus, there is a gap in knowledge regarding the oncogenic role of NRAS in the progression of CRC and factors driving resistance to standard-of-care drug therapy.

Our interdisciplinary team identified a patient with exceptionally aggressive, chemorefractory CRC metastatic to the liver, lungs, peritoneum, and brain (>100 individual metastatic sites). Clinical NGS testing (8) for an actionable CRC gene panel identified a novel *NRAS* internal tandem duplication (ITD) of 10 amino acids within the switch II domain. The prevalence of such *RAS* ITDs in CRC is unknown, and at the time of initial diagnosis (2015), we found literature reports of only three prior CRC patients with this type of duplication in *KRAS* (2,9,10). Subsequently, other reports of *RAS* ITDs have emerged including one report of a pediatric patient with a hematologic malignancy(11) and a report of *RAS*-family ITDs in up to 2% of vascular malformation/overgrowth syndrome cases (12).

We hypothesized that this previously uncharacterized *NRAS* ITD was the primary driver of this aggressive case of CRC. With written consent, the patient’s tumors were harvested for research after his death; we proceeded with whole exome DNAseq of the primary tumor and metastases from the lung and brain to identify other potentially pathogenic somatic mutations that drove the evolution of the patient’s disease. We also re-processed the TCGA CRC datasets (13) with our bioinformatic algorithms (14) to determine whether this type of *RAS* ITD was under-called in prior analyses. Finally, we performed extensive functional and structural characterization of this ITD mutation to determine its effects on RAS biology.

## Results

### Clinical course of metastatic, chemo-resistant CRC

A 48-year-old male presented with abdominal pain and iron-deficiency anemia. Colonoscopy was performed and revealed a large bleeding mass at the rectosigmoid junction, diagnosed as invasive adenocarcinoma on biopsy. Staging evaluation detected metastatic regional lymphadenopathy, bilateral pulmonary nodules, and liver lesions consistent with metastasis (stage IV disease). The primary tumor was resected because of tumor perforation of the GI tract. Initial *KRAS* Sanger sequencing (the institutional standard of care at that time) was negative for pathogenic mutations. After post-operative recovery, the patient initiated palliative chemotherapy with the mFOLFOX-6 regimen. He received 12 cycles of therapy and experienced a favorable radiologic response. He then received six months of maintenance oral capecitabine treatment; two months into this course, he was diagnosed with metachronous chronic myelogenous leukemia (CML) for which he received imatinib therapy concurrent with continued capecitabine. Six months after initiation of capecitabine, he developed systemic progression of disease with new and progressing lung and liver nodules, as well as a new cerebellar metastasis. Right suboccipital craniotomy was performed to resect the cerebellar tumor. This tumor was confirmed to be metastatic colorectal adenocarcinoma on histopathology. Palliative second-line FOLFIRI therapy was given, but the patient experienced rapid progression of systemic disease after only four cycles. Re-challenge with FOLFOX with addition of bevacizumab was tried, but the patient developed an adverse reaction to oxaliplatin. Next-line regorafenib was administered, but within four weeks the patient developed further rapid systemic progression and two additional cerebellar metastases. After stereotactic radiosurgery, next-line therapy was attempted with the EGFR inhibitor panitumumab because previous testing had indicated the tumor was *KRAS* wildtype. However, he again developed clinical and radiographic progression within six weeks.

### Next-generation sequencing (NGS) identifies a novel *NRAS* ITD in cerebellar metastasis

Because the patient’s cerebellar metastases progressed on panitumumab, a focused clinical NGS panel was used to assess the cerebellar lesion for mutations in *KRAS, NRAS, HRAS, BRAF*, and *PIK3CA*. NGS identified a novel in-frame duplication of 30 nucleotides in exon 3 of *NRAS* (NM_002524:c.164_193dup, p.I55_Y64dup). At the time of this diagnosis, no in-frame insertions of this size or larger were catalogued within *NRAS* exon 3 in the COSMIC database (15); thus, any potential clinical or biologic significance of this alteration was initially unclear. This *NRAS* ITD is located within the switch II domain of the protein, which includes the Q61 amino acid critical for GTP hydrolysis (16); thus, we inferred this mutation might prevent GAP-mediated inactivation of the NRAS protein and lead to constitutive downstream signaling. Given the paucity of clinical information but plausible biologic mechanism, this mutation was interpreted as a likely pathogenic variant with tier 2 clinical significance (17) for prediction of resistance to anti-EGFR therapies. No other mutations in *BRAF, PIK3CA, KRAS, or HRAS* were identified. The patient acutely developed worsening fatigue, nausea, and vomiting; he was diagnosed with subdural bleeding secondary to further progression of brain metastases. He and his family elected for hospice care, and he passed away two weeks later.

### Post-mortem examination and focused NGS testing of multiple tumor sites

An autopsy was performed, which catalogued >100 individual metastatic lesions within the bilateral lungs, multiple lymph node chains, liver (including a dominant 12.0 × 10.0 × 4.5 cm lesion), pancreas, gallbladder, thyroid, bilateral adrenal glands, and brain parenchyma (left insula and right inferior cerebellum). No residual tumor was identified at the surgical site of the primary tumor. To further characterize the patient’s malignancy, three metastatic sites (cerebellum, lung, and liver) plus the archived primary tumor from the prior surgical resection were assessed by a clinically-focused solid tumor NGS panel of 13 genes. The *NRAS* ITD was identified in all four samples, confirming that this mutation was present in the primary tumor and in three independent metastatic sites that developed at different times during the patient’s disease course. In addition, a pathogenic *TP53* mutation (NM_000546: c.824G>T, p.C275F) was identified in the primary tumor and all three metastatic sites. No other pathogenic mutations were identified in the remaining 11 clinically-relevant genes assessed in these four samples at the time of autopsy.

### Exome sequencing of primary and metastatic tumors

Given the lack of additional potential driver mutations on the targeted clinical cancer NGS panel, we used whole exome NGS of the archived primary tumor, lung metastasis, and cerebellar metastasis to more extensively explore whether other genomic drivers were responsible for the aggressive clinical course of the disease. Variant annotation and filtering identified 847 potentially significant somatic variants in the primary tumor and an additional 551 potentially significant variants that were unique to the two metastatic lesions (i.e., variants not identified in the primary tumor) (Figure 1A). These variant lists were further prioritized for loss of function variants (frameshift indels or nonsense), in-frame indels, canonical splice site variants, and single nucleotide missense variants with high-to-moderate Mutation Assessor functional impact (18) scores; this process left 169, 58, and 72 prioritized variants for review in the primary, lung, and central nervous system (CNS) lesions, respectively (Supplemental Data File 1).

**Figure 1:**
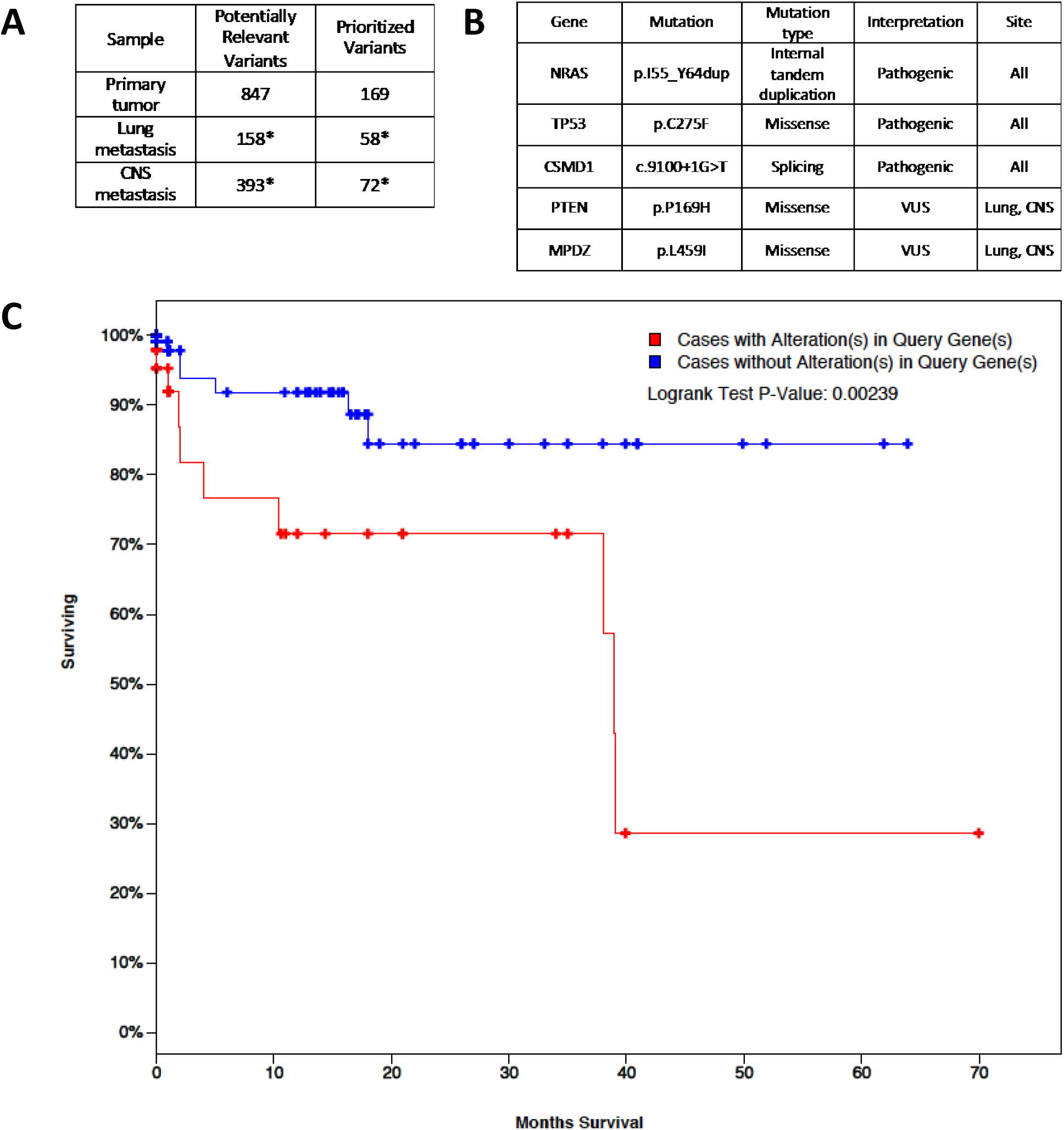
Whole exome sequencing of a clinically aggressive CRC suggests the NRAS-ITD is the primary oncogenic driver mutation. A) Overview of filtered variants prioritized for pathology review in whole exome NGS of the patient’s primary tumor, lung, and CNS metastases; * indicates variants unique to the metastases vs. the primary tumor. B) Clinically relevant variants, including tumor sites where identified. C) Overall survival of patients with NRAS or CSMD1 mutations (red, N=48, median OS 38.9 months) was significantly (p=0.002) shorter than patients without mutations in these genes (blue, N=162) in the CRC TCGA cohort.

The *NRAS* ITD and pathogenic *TP53 p.C275F* missense mutations were verified by DNA exome analysis in all three specimens (Figure 1B). In addition, a splice site mutation was identified in all tested tumor sites that abolished the splice donor site of exon 58 in *CSMD1* (which contains 70 total exons). CSMD1 (CUB and Sushi Multiple Domains 1) is a transmembrane protein that is not well-characterized functionally, but studies indicate it acts as a tumor suppressor in multiple tumor types (19–23). Specifically, mutations of *CSMD1* appear to be enriched in stage 3 and 4 colorectal carcinomas (20), and loss of *CSMD1* function by both genomic and epigenomic mechanisms has been linked to earlier age of onset (21). Thus, it is likely this mutation shared across all tumor sites played a pathogenic role in the aggressive phenotype of the tumor.

Two mutations that appeared limited to the CNS and lung metastases were identified; both were classified as variants of uncertain, but possible, significance. The p.P169H missense mutation in *PTEN* occurs in the TI catalytic loop, which is a hotspot for missense mutation, but *in vitro* functional studies indicate this specific variant has nearly wild-type levels of PIP phosphatase activity (24). Thus, it is unclear whether this mutation would have affected PI3K signaling in our patient’s metastatic lesions. The second mutation of interest restricted to the metastases was a p.L459I missense mutation in *MPDZ* (multiple PDZ containing protein). No specific information on this variant could be identified in somatic mutation databases or the published literature. Nonetheless, loss of MPDZ function has been associated with metastasis and poor prognosis in breast cancer (25); thus, the identification of this mutation in only metastatic sites was intriguing, but not sufficiently conclusive of a pathogenic role.

To gain a broader perspective of the mutation spectrum in this patient’s cancer, we performed functional pathway analysis (26,27) of the genes that harbored the 299 total prioritized mutations. No individual pathways were statistically enriched in either the primary, lung, or CNS samples. The most highly represented pathways (Supplemental Figure 1) included angiogenesis, WNT signaling, p53 regulation, and other cell signaling pathways. Mutated genes restricted to the metastases revealed accumulation of additional alterations in similar pathways prevalent in the primary tumor (i.e., angiogenesis, WNT and cadherin signaling), with the addition of inflammatory cytokine signaling and the cholecystokinin receptor pathway, which has been associated with angiogenesis, cell survival, and invasion (13).

Overall, somatic exome analysis of the patient’s primary colon tumor, lung, and CNS metastasis indicated that the *NRAS* ITD was likely the most significant driver mutation in this patient’s malignancy. Loss of tumor suppressive functions of *TP53* and *CSMD1* likely contributed to the aggressive phenotype of the malignancy. Interestingly, *NRAS* and *CSMD1* mutations were associated with decreased overall survival in the TCGA (13) CRC dataset (Figure 1C).

### Interrogation of TCGA datasets to assess prevalence of *RAS* ITDs in colorectal cancer

The original analysis of the TCGA colorectal adenocarcinoma project (13) did not report any similar insertion mutations within *RAS* genes. We hypothesized that this type of duplication may have been missed by the bioinformatics pipelines that produced the published datasets. Therefore, we performed a focused re-analysis of the *RAS* genes using the primary sequencing files from 634 patient samples in the TCGA cohort with the ScanIndel bioinformatics pipeline (14), which has been optimized to detect longer insertion mutations. No clinically significant in-frame insertion mutations within *NRAS, KRAS*, or *HRAS* were identified in that dataset. This analysis suggests that somatic in-frame insertion mutations (including ITDs) within *RAS* genes are likely a rare event in colorectal adenocarcinoma development.

### Cell-based functional characterization of NRAS ITD confirms constitutive activity

To determine the functional activity of the NRAS ITD protein, we used bioluminescence resonance energy transfer (BRET) to detect protein-protein interactions in live, intact cells(28,29). We co-transfected HEK293T cells with plasmids expressing mutant or wildtype RAS proteins attached to the HaloTag acceptor construct with plasmids expressing RAF1 tagged with the NanoLuc bioluminescent donor construct. In saturation experiments, cells were transfected with a constant amount of the RAF1-NLuc donor construct in the presence of increasing amounts of the indicated RAS-HaloTag acceptor constructs. In 12-well plates, we generated saturation curves for the interactions between RAF1 and wild-type KRAS, wild-type NRAS, ITD mutant KRAS, Q61R mutant KRAS, and ITD mutant NRAS (Figure 2). Curves were produced by a hyperbolic linear regression and the BRET_50_ was calculated. The BRET_50_ is the transfected DNA mass of acceptor which generates a bioluminescence signal at 50% of maximum; lower values are consistent with higher affinity of interaction. These experiments showed more affinity between the internal duplication mutant interactions with RAF1 versus the wild-type proteins (Figure 2B, 2C). Biologically, other known carcinogenic point mutations (KRAS Q61R) recruit RAF1 with higher affinity, leading to more downstream signaling (Figure 2A). These results suggest that the novel 10 amino acid ITD (in both NRAS and KRAS) drives interactions with the effector RAF1 at a similar level of potency to the well-characterized Q61R oncogenic missense mutation.

**Figure 2:**
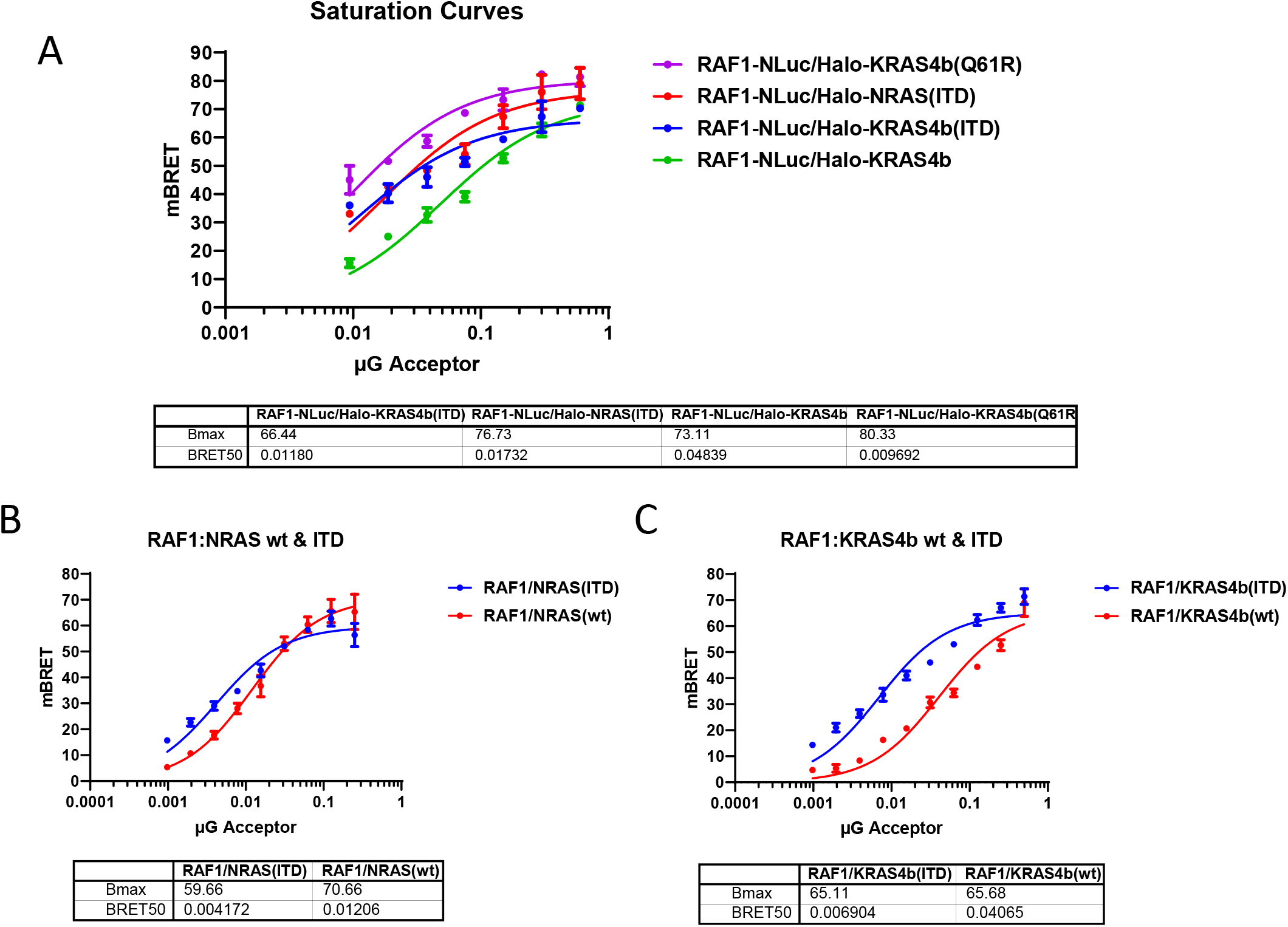
Switch II Internal Tandem Duplication of both NRAS and KRAS proteins increases the affinity for RAF effector. A) mBRET saturation curves comparing KRAS wildtype (Kras4b), KRAS Q61R missense mutant (4b Q61R), KRAS ITD (4b ITD), and NRAS ITD. B,C) mBRET saturation curves comparing wildtype vs. ITD mutant forms of NRAS and KRAS, respectively.

### Downstream MAPK signaling is activated by the RAS ITD

To further demonstrate biologic relevance of the increased affinity of NRAS ITD to RAF in cells, we evaluated the effect of expression of the internal duplication mutants on downstream signaling pathways. Transfection of increasing amounts of both NRAS and KRAS ITD mutants into HEK293T cells showed a significant increase in MAP Kinase signaling as compared to the wild-type proteins, measured by both ERK and MEK phosphorylation in Western blots (Figure 3A-B). When compared to the oncogenic mutant KRAS G12D, the KRAS internal tandem duplication mutant showed a similar increase in both ERK and MEK phosphorylation relative to protein expression (Figure 3C). In contrast, transfection of either ITD or G12D mutant RAS did not have an appreciable effect on AKT phosphorylation (Figure 3A-C); we did not further pursue this observed lack of PI3K-AKT pathway response in cell lines other than HEK293T, and suggest that other RAS-independent mechanisms may sustain AKT phosphorylation in this cell line (30,31). These results were consistently produced across three independent experiments, providing clear evidence that a functional consequence of the ITD mutation is constitutive activation of the oncogenic MAP kinase signaling pathway.

**Figure 3.**
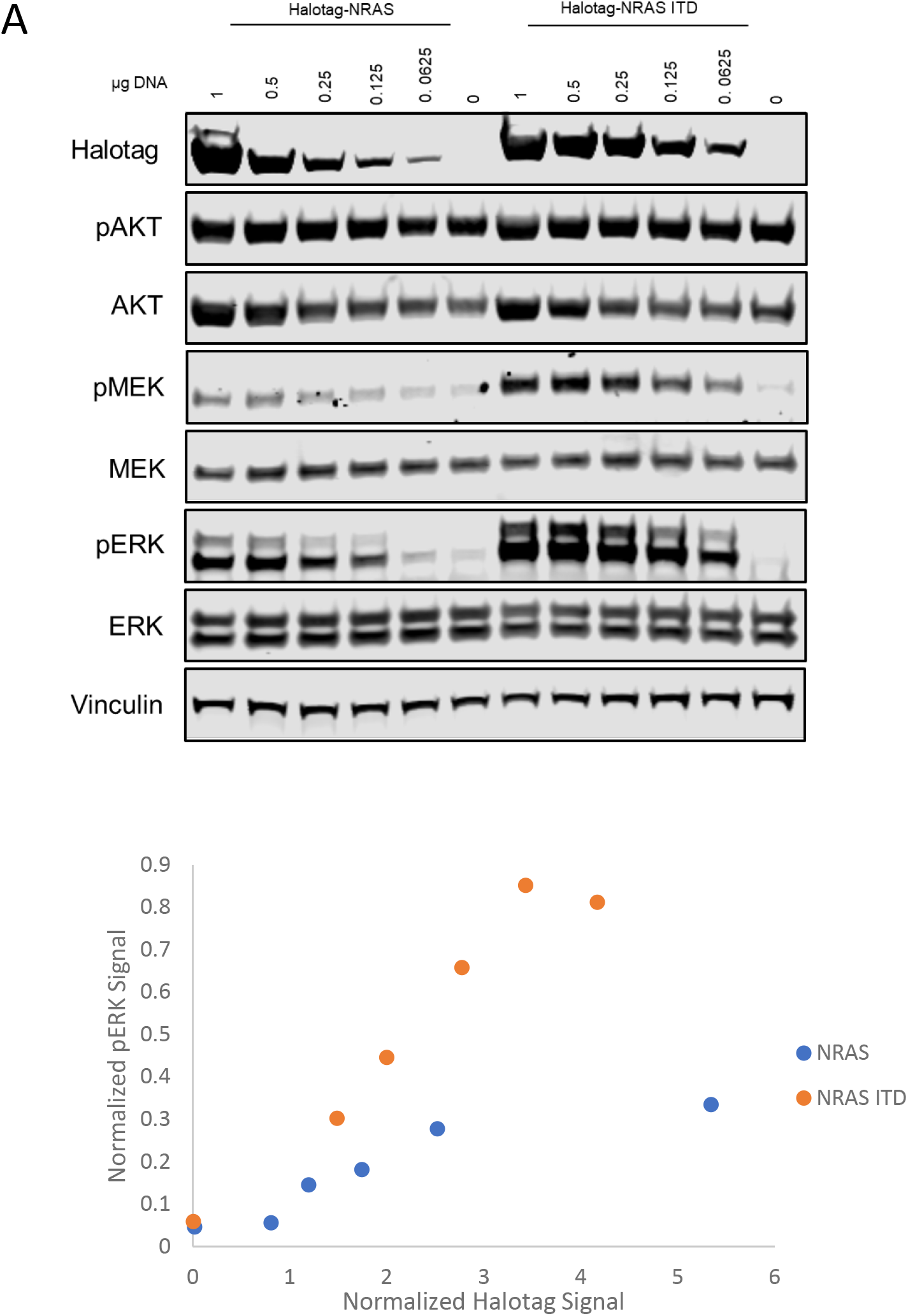
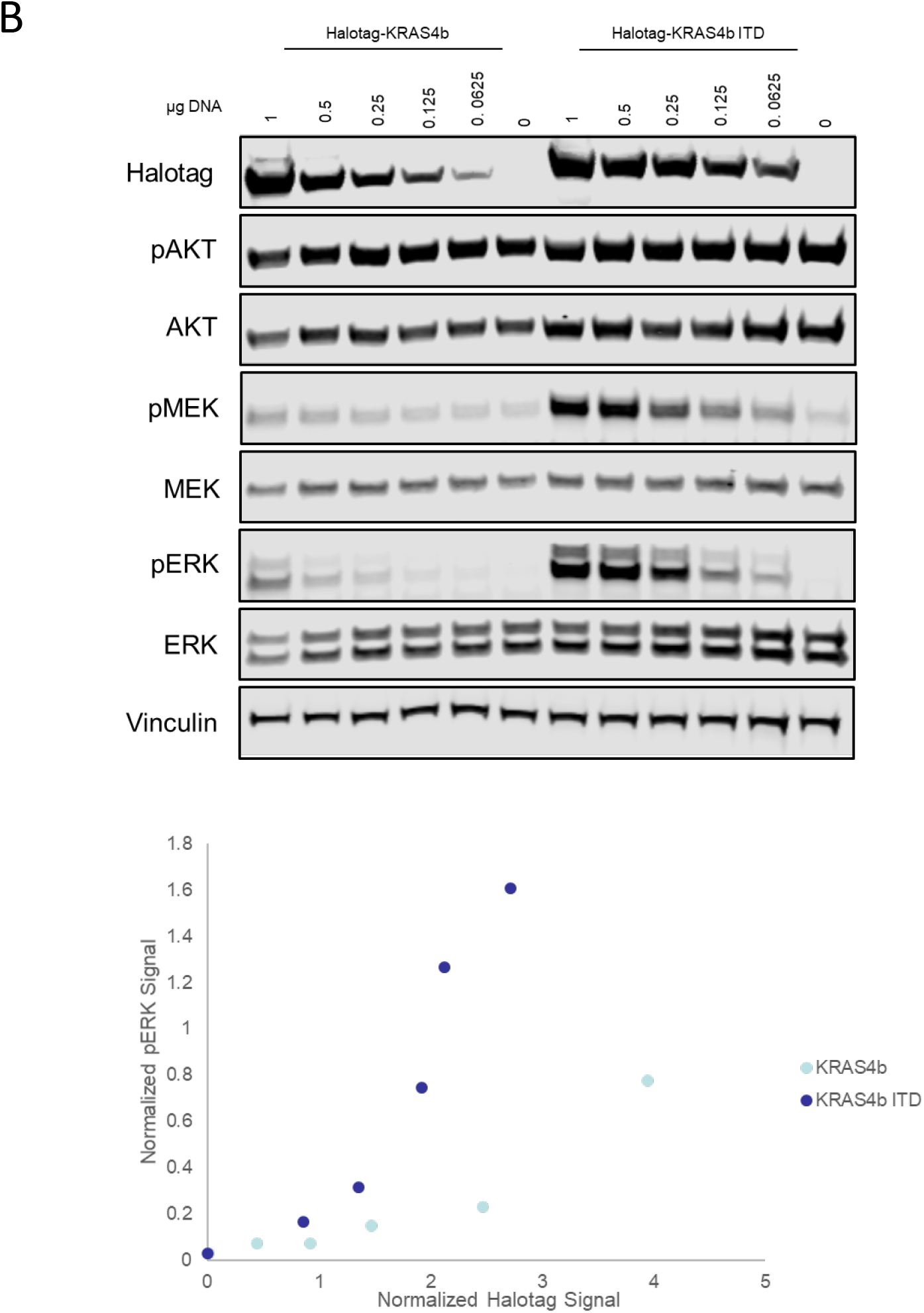
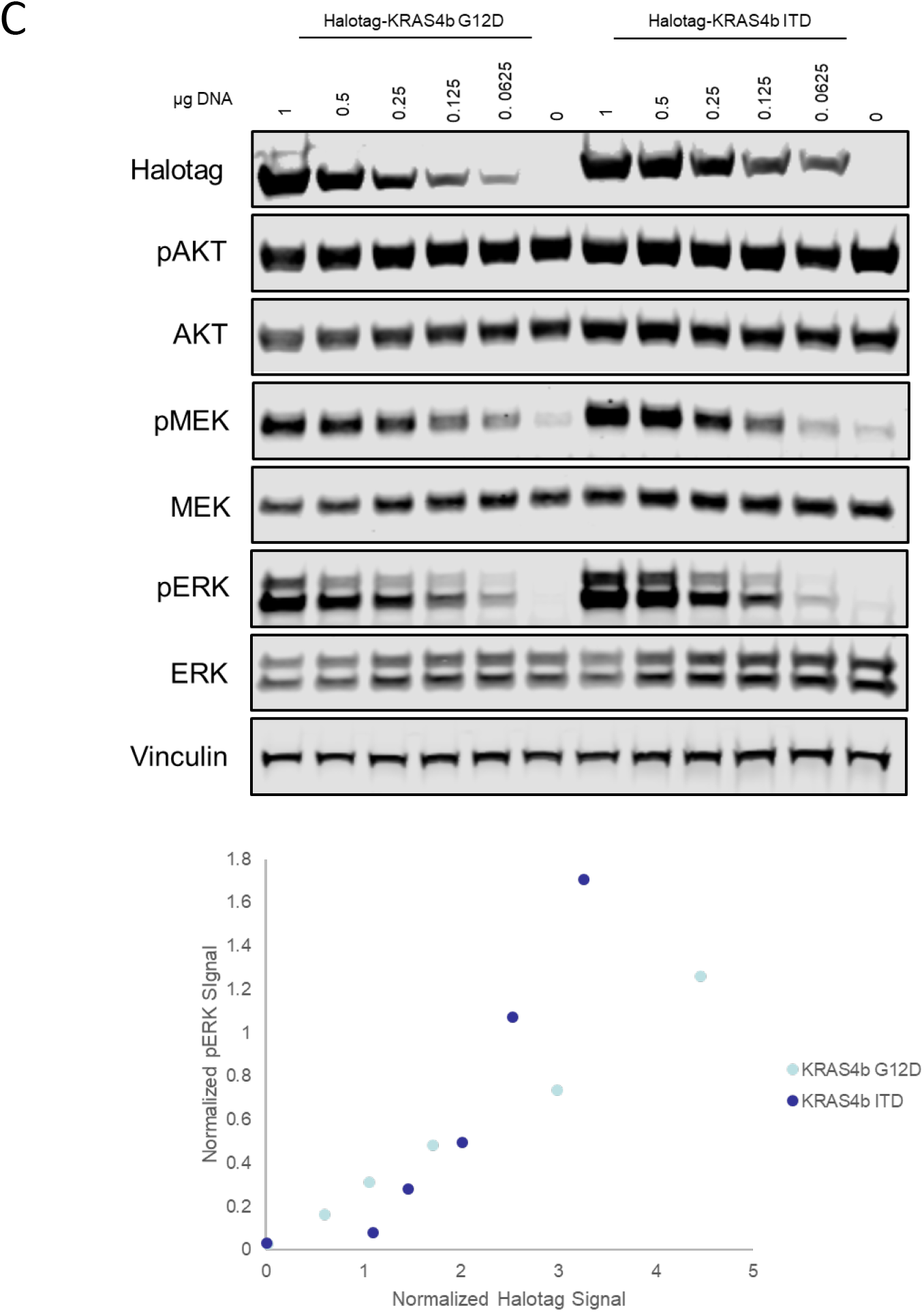
Expression of ITD mutant RAS leads to MEK and ERK activation. A) Comparison of phospho-MEK, ERK, and AKT levels after wt vs. ITD NRAS transfection. B) Comparison of wt vs. ITD KRAS transfection. C) Comparison of G12D and ITD KRAS mutants. Each protein was expressed with a Halotag fused to the C-terminus, so that increases in MAPK activity could be normalized to expression levels via Western blot.

### Recombinant RAS ITD binds to RAF1 but not to NF1 (neurofibromin 1) GAP

To examine the effect of the ITD in RAS on its interaction with GAP and its effector proteins, we purified the recombinant GTPase domain (1–169) of KRAS with and without the ITD and carried out binding studies using isothermal titration calorimetry (ITC). Due to ease of expression, purification, and crystallization, we used recombinant KRAS4b (GTPase domain) for biochemical and structural studies. This approach to understanding the structural consequence of the ITD is reasonable because RAS isoforms (*NRAS, KRAS* and *HRAS*) share 100% sequence identity in the N-terminus of the G domain (aa 1–86) and 82% sequence similarity within residues 87–169. We compared the binding of wild-type KRAS and KRAS ITD with NF1 GAP and RAF1 effector protein. ITC experiments showed that WT KRAS bound to NF1 (GRD; GAP-related domain) with high affinity (Kd = 1 uM), whereas KRAS ITD showed complete loss of binding to NF1, suggesting that the ITD resulted in fully impaired GAP-mediated GTPase activity (Fig. 4A, upper panel). Unlike NF1 GAP, both WT KRAS and KRAS ITD bound to RAF1-RBD with similar affinity *in vitro*, confirming that the ITD does not negatively affect RAF1 (RBD; RAS-binding domain) binding (Fig. 4A, lower panel). This biochemical evidence is consistent with the cell biology data (Figures 2 and 3) showing increased physiologic interaction between RAS-ITD and RAF with subsequently increased MAPK signaling.

**Figure 4:**
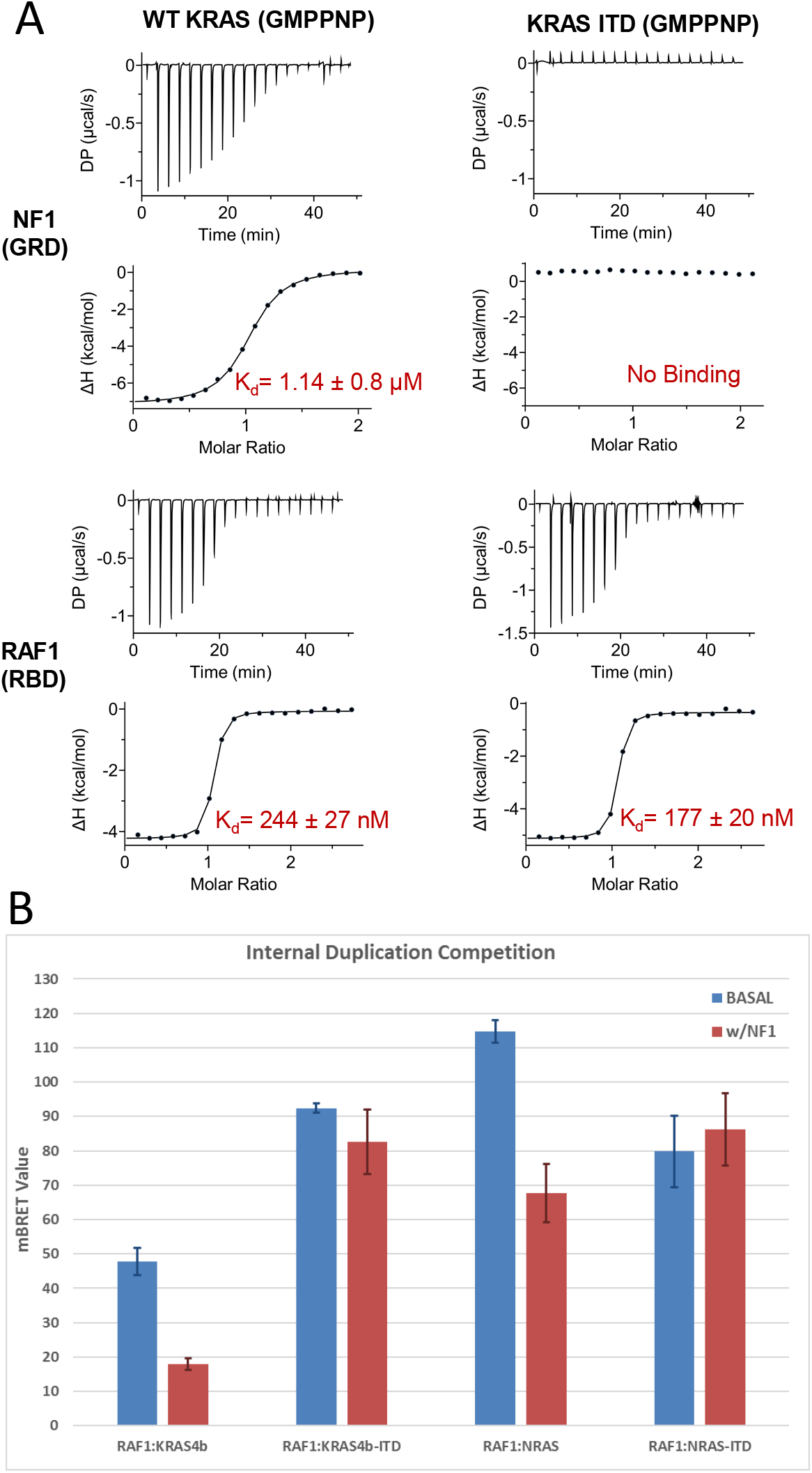
The ITD mutation in RAS does not affect RAS-RAF1 interaction but blocks RAS-RasGAP binding. A) Isothermal titration calorimetry experiments to measure the dissociation constant for GMPPNP-bound WT KRAS and KRAS ITD with RasGAP NF1 (GRD) and effector RAF1 (RBD). Differential power (DP) is a measure of energy required to maintain isothermal conditions between the reference cell and the sample cell. B) mBRET saturation values assessing the ability of NF1-GAP co-expression to squelch RAF-RAS interaction for wild-type and ITD mutant KRAS and NRAS.

To explore the lack of NF1-GAP:RAS-ITD binding further, we returned to our BRET cell-based expression system. By co-expressing NF1 with the RAF1 and RAS constructs, we demonstrated biologically that NF1 expression significantly decreased the peak mBRET signal for wild-type RAS constructs, but had no effect on the RAS-ITD constructs (Fig. 4B). This finding confirmed within an intact cell model system that the switch II ITD prevented effective interaction of RAS with GAP protein, leading to sustained activity.

### Crystal structure of RAS ITD provides rationale for its oncogenicity

To understand the effect of ITD on RAS tertiary structure and RAS interactions with GAP and effector proteins, we crystallized and solved the structure of GDP-bound KRAS with ITD to a resolution of 1.85 Ang (Fig. 5A and Supplemental Table 1). In this structure, we observed a large conformational change in the switch II region because of the ITD. The first seven amino acids in the ITD formed a loop in the middle of the switch II region, which protruded away from the core of the protein. To accommodate ten amino acids inside the switch II, the helix alpha2 transitioned into a loop structure where additional residues due to the ITD and six wild-type residues from Gly75 to Cys80 formed a long loop. Because of the structural mobility of this loop, we observed no electron density for the last three residues of the ITD and the six residues present at the end of switch II (amino acids 75-80) likely because of the multiple conformations of these residues inside the loop. Structural comparison of GDP-bound KRAS-ITD with GDP- and GMPPNP-bound wild-type NRAS highlighted the difference in the switch II because of the ITD, and shows similar tertiary structure including the switch I region (Fig. 5B and 5C).

**Figure 5:**
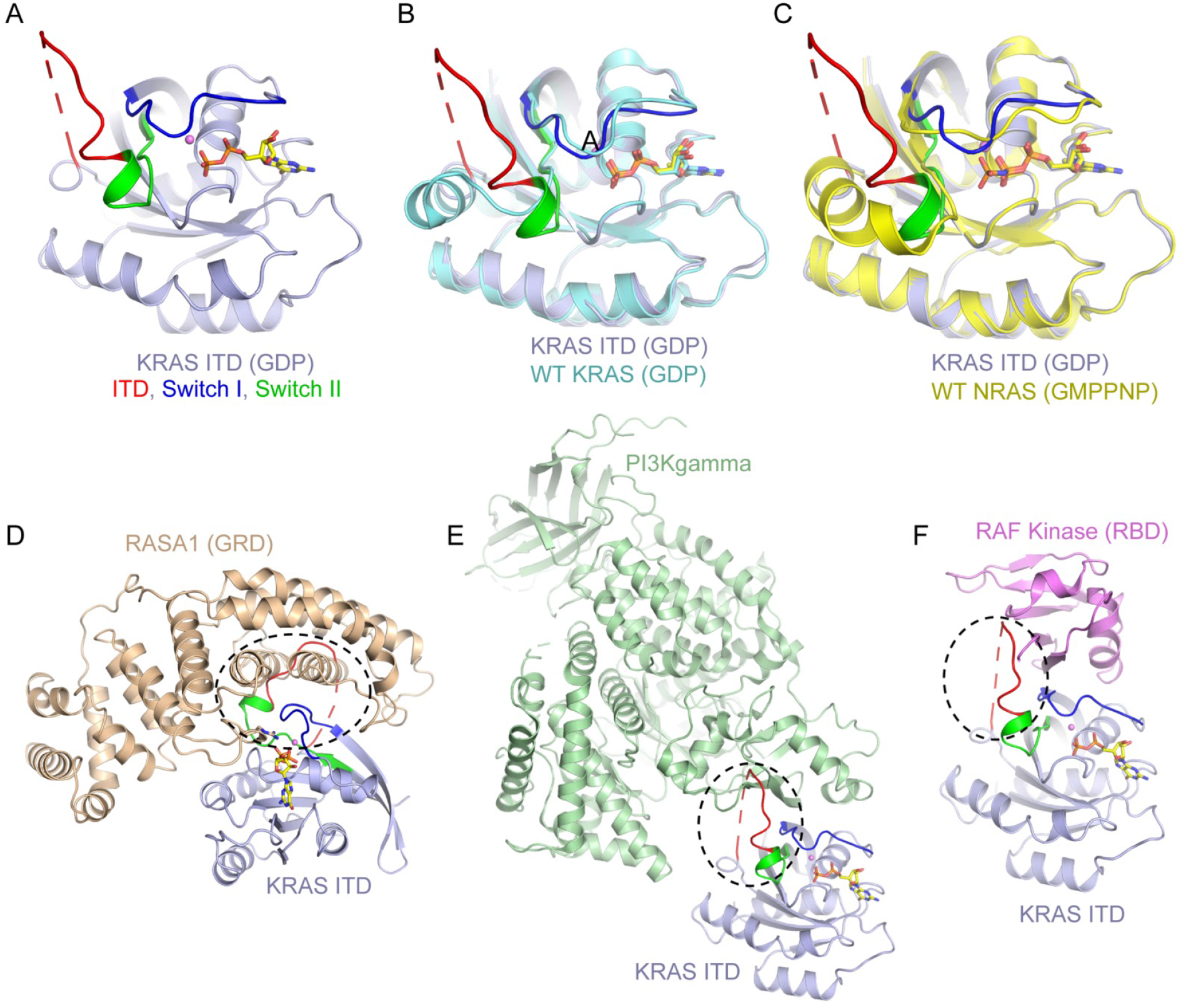
Crystal Structure of GDP-bound KRAS ITD and insights into the effect of ITD on RAS interaction with GAP and effector proteins. **A)** The tertiary structure of GDP-bound KRAS-ITD. **B, C)** Structural superposition of GDP-bound KRAS ITD with GDP-bound WT KRAS and (C) GMPPNP-bound WT NRAS. **D, E, F)** Models of KRAS ITD in complex with (D) RASA1 (GRD), (E) PI3Kgamma, and (F) RAF1 (RBD) generated using the structural superposition of KRAS ITD on HRAS present in HRAS-RASA1, HRAS-PI3Kgamma, and HRAS-RAF1 (RBD) complexes. These models suggest that the ten amino acid insertion (shown in red) in KRAS would sterically clash with RASA1 GAP and PI3Kgamma and not with RAF1. The region where ITD residues interact with GAP and effector proteins are highlighted in dotted circle colored in black.

Structural superposition of the KRAS ITD with the HRAS-RASA1 complex, HRAS-PI3Kgamma complex, and HRAS-RAF1 complex showed a steric clash of RAS ITD in the case of RASA1 and PI3Kgamma, but not in the case of RAF1 (Fig. 5D, 5E and 5F). This observation is aligned with previous results that have shown the RAS-RAF1 interaction is limited to switch I (32), whereas RASA1 and PI3Kgamma interact with RAS protein via both switch I and II regions. Unlike oncogenic mutations that are limited to a different side chain at a single amino acid, this novel ITD of 10 residues creates a significant steric clash that is likely to result in complete loss of binding of GAP protein to RAS ITD.

## Discussion

Identification of novel isoforms of oncogenic *RAS* over the past decade has expanded the pool of testable variants that help predict biological behavior as well as serve as potential predictive biomarkers of therapeutic efficacy of drugs such as inhibitors of EGFR in patients with metastatic CRC. Despite expansion of *RAS* sequencing platforms and greater capability for having this clinical testing performed more widely, there remains concern that additional variants remain undetected, and can thus affect clinical outcomes. Here, we identify a novel *NRAS* ITD that disrupts the switch II domain and prevents GAP-mediated GTP hydrolysis. Extensive exome analysis did not reveal any other activating, driver mutations. The NRAS ITD was found in the context of loss-of-function alterations in the tumor suppressors TP53 and CSMD1. This *NRAS* ITD was the only clear oncogenic driver (by whole exome DNAseq) in an especially aggressive and therapeutically resistant form of CRC. Through interrogation of the TCGA database, we found that ITDs have a low prevalence in CRC. Its biologic activity is nonetheless substantial, as this ITD facilitates RAS-RAF interaction in a cell biology system that leads to increased MAPK signaling, as we demonstrate here using in vitro analysis. More specifically, the ITD blocks the interaction of RAS-GAP in vitro, and our cell biology system confirms that GAP co-expression does not diminish sustained RAS-ITD:RAF1 affinity/interaction. Furthermore, by solving the crystal structure of GDP-bound RAS-ITD, we provide convincing structural evidence that the 10 amino acid ITD blocks RAS-GAP binding, and provide insights into the interactions of the RAS-ITD with other effector proteins. Our data suggest that ITDs of this length lead to constitutive activation of RAS with significant biologic and clinical consequences.

The work shown here is consistent with findings seen in other limited reports of RAS ITDs. ITDs in general are indeed rare in somatic malignancies, but functional and biochemical data are consistent with an oncogenic phenotype (11). Eijkelenboom et al. have reported that RAS ITDs (particularly involving HRAS) are slightly more common in vascular malformation/overgrowth syndromes (2% prevalence) (12). As the oncogenic and other biologic properties of RAS isoforms (*KRAS* vs. *NRAS* vs. *HRAS*) differ, we likewise postulate that the clinical effects of their respective ITDs may vary as well between RAS family members. Further, the size range of previously reported ITDs is 5-10 amino acids (11). The accumulating literature on RAS ITDs indicates this structural variability may drive variability of biochemical function: differences in nucleotide exchange rates, effector interactions, GTPase activity, and likely other downstream effects suggests that the specific size of the duplication/in-frame insertion within this region is critically important to defining downstream biologic consequences.

Exome sequencing of the archived primary tumor as well as anatomically distinct metastatic deposits that developed at different points in the disease course confirmed that three pathogenic mutations (*NRAS ITD, TP53 p.C275F*, and *CSMD1* splice site/loss of function) were conserved between all sites, consistent with a driver role in the disease. Although additional somatic variants were identified in genes with clear roles in the pathogenesis of colorectal carcinoma (such as *CTNB1, DDR2, PTEN*), the specific variants found in this patient’s cancer did not appear to be consistent with clearly pathogenic roles based on the type of alteration (*CTNB1, DD*R2) or existing data from laboratory models (*PTEN*). Analysis of the variants restricted to the metastatic sites did not identify an additional set of clearly pathogenic driver mutations. Nevertheless, this analysis did identify a missense mutation in *MPDZ* (multiple PDZ containing protein), which is involved in cell-cell adhesion. Loss of *MPDZ* function is associated with higher rates of metastasis (25), so this finding could represent a possible mechanism promoting the extensive metastatic dissemination of this patient’s disease. However, because information on the specific missense variant identified (p.L459I) was lacking, it could not be definitively classified as a pathogenic mutation in this case. This analysis suggests the combination of *NRAS, TP53*, and *CSMD1* mutations was the primary driver of this aggressive colorectal carcinoma and demonstrates the complexity of clinically interpreting a single patient’s tumor exome data to better understand tumor behavior on a personalized level.

Why are our data important to the field of RAS biology? Few cases of RAS ITDs have been published to date. In one study from Poland, 163 colon adenocarcinomas were examined using Sanger sequencing of *RAS* exons 2-4 (10). One case with a *KRAS* insertion was identified in an early-stage CRC (T2N0M0) with low level of microsatellite instability (MSI). This particular patient was considered cancer-free at 43 months before being lost to follow-up (10). White et al. also identified and characterized a novel 7 amino acid *KRAS* ITD in a pediatric patient with juvenile myelomonocytic leukemia (JMML)(11). Additionally, Guedes et al. identified two samples with *KRAS* duplications in their study of metastatic colorectal cancer, but did not further elaborate on the clinical characteristics of these cases (9). As compared to the reports from White et al.(11) and Eijkelenboom et al. (12), our data suggest more robust downstream activation of MAPK by the 10 amino acid duplication in both KRAS and NRAS. We also provide data gathered from in-depth *in vitro* evaluation of ITD activity, and additionally describe the first crystal structure of the RAS ITD. Elucidation of the stream of mutations and similarities vs. differences of the *NRAS* ITD and other mutations between the primary and metastatic tumors helped to confirm the *NRAS* ITD as the primary oncogenic driver in this case, whereas *MPDZ* for example is a likely secondary mutation as it was present in the metastatic clones but not identified in the primary tumor itself. There are a number of lessons that can be garnered from the latter, lessons that are similar to published reports of *RAS* driver and other passenger mutations in similar RAS-driven cancers, including most prominently pancreatic cancer (33).

Taken together, this clinic-to-lab endeavor represents a process that effectively identified a clinically challenging case for which a potential solution was uncovered through rigorous molecular and biochemical analysis. Had this *NRAS* ITD been characterized prior to the patient’s diagnosis or at any point during his treatment, EGFR inhibitor therapy would not have been administered even in the context of wild-type KRAS expression because of the lack of an anticipated response (34). Although the ITD subclass of *RAS* alterations is still relatively rare, our finding, in combination with those of the groups listed above, indicates that RAS ITDs have important clinical impact and warrant further biologic study to understand the context-specific impact on cancer progression and therapeutic response.

In addition, the clinical and scientific implications of this study are that acquisition and utilization of tumor and non-malignant tissue specimens from a research autopsy circumvented concerns of tumor heterogeneity affecting our results, via sample bias occurring by obtaining only a small portion of a resected tumor for research purposes. In fact, the strategy of rapid research autopsies performed immediately post-mortem has gained increasing ground in the past few years as a method to overcome concerns of tissue breakdown as well as having sufficient tissue available for full analyses (35). The issue of accurate, comprehensive investigation of tumor genomics is not only of paramount concern for the field of tumor biology and molecular oncology in general, but it also resonated with this specific individual case. In this index patient, initial focused *KRAS* testing was negative and guided therapy on a *RAS* wild-type algorithm. The chronology of the case straddled the advent and incorporation of comprehensive RAS sequencing at our and many other institutions. It was only upon further repeat testing, following failure of the patient’s metastatic tumors to respond to EGFR inhibition, that the novel *NRAS* ITD was identified. Insights from this case caution us to continually evaluate whether current standards of testing are sensitive enough to detect all clinically-relevant *RAS* alterations, however rare they might be, as they may have clinical implications for patient prognosis.

Potential limitations of this study include the fact that these extensive exome results from multiple sites of disease represent the underlying genomics of one patient, and thus cannot readily be extrapolated to all *NRAS*-driven CRC tumors. Also, there is no certain way to distinguish what effects, if any, the concurrent diagnosis of CML had on the biologic activity of the aggressive course of the patient’s CRC in general. The mutation profile of *NRAS* is different from that of *KRAS*; therefore, while they are structurally conserved, the small differences in structure between these RAS subtypes likely manifest as diverse functions, and the crystal structure of the *KRAS* ITD may not necessarily reflect its counterpart in *NRAS*. Our functional in vitro studies were performed in HEK293 cells to provide proof-of-concept analysis. Further work in cells derived from colonic epithelium could provide further confirmation of the effect of the *NRAS* ITD mutation on biochemical and biological properties in CRC specifically. *In vitro* analysis involved overexpression of the ITD; the physiologically relevant expression levels are unknown but could be explored in further studies.

In summary, we have identified a novel *NRAS* ITD and confirmed its activity as an oncogenic driver associated with biologically aggressive metastatic CRC, as it was conserved between primary and metastatic tumors in vivo, had constitutive biologic activity, and a novel crystal structure that identifies it as a distinct and new form of mutant *NRAS*. Added to a growing list of new oncogenic alterations of *RAS*, our data provide impetus for continuous quality improvement in pan-*RAS* molecular analysis to ensure our ability to identify these alterations adequately in the clinical setting.

## Materials and Methods

### Patient samples

A clinical research autopsy was performed, and tumor samples were snap frozen or stored in formalin fixed, paraffin embedded (FFPE) blocks.

### Targeted Next Generation Sequencing

An amplicon-based target enrichment of portions of 13 genes (Supplemental Table 2) followed by next generation sequencing (NGS) was performed on all specimens using a CLIA-validated workflow (8). A subset of this panel is reported clinically for colorectal carcinoma (*KRAS, NRAS, BRAF, PIK3CA, HRAS*). DNA was extracted from formalin fixed, paraffin embedded (FFPE) samples using the QIAamp DNA mini FFPE tissue kit and deparaffinization solution (Qiagen, Hilden, Germany) and quantified using a Qubit 2.0 fluorometer (ThermoFisher Scientific, Waltham, MA). Amplicon enrichment was performed using a custom designed primer set on the Fluidigm (San Francisco, CA) Biomark Access Array system followed by sequencing with 2×225 base pair reads on a MiSeq instrument using version 3 chemistry (Illumina, San Diego, CA). FASTQ files were processed through a custom bioinformatics pipeline (ScanIndel) for mapping, indel realignment, and variant calling (14).

### Exome Next Generation Sequencing

Four samples (3 fresh frozen, 1 FFPE) were prepared for whole exome NGS: metastatic lung, metastatic CNS, archived primary colon adenocarcinoma, and normal tissue. Normal tissue was collected for assessment of germline variation; the patient had concurrent chronic myelogenous leukemia, so peripheral blood could not be used. DNA from snap frozen tissues were extracted using the DNeasy Blood & Tissue Kit (Qiagen, Hilden, Germany). Libraries were enriched using SureSelect QXT reagents and V6+COSMIC exome hybrid capture baits (Agilent, Santa Clara, CA). Sequencing was performed with 2×125 base pair reads on a HiSeq 2500 with V4 chemistry (Illumina, San Diego, CA). FASTQ files were processed as described above. Average coverage was >200X for all four samples. All variants with variant allele fractions >0.3 in normal tissue were considered germline. To identify somatic alterations in tumors, these germline variants were filtered out of the tumor sample variant call files. To identify unique somatic variants that either arose in metastatic tumors or were very rare subclones not detected in primary tumor, all variants identified in the somatic primary colon variant call file were then further filtered out of the somatic variant call files of the lung and CNS metastases.

Variant call files were annotated using Cravat (36,37) and Annovar (38). Annotated variant files were filtered to remove synonymous variants not affecting splice sites and any passenger somatic variants that had been recorded as polymorphisms in healthy human population databases with minor allele frequencies > 0.01. Variants were priority filtered for all stopgain, frameshift, in-frame indel, and canonical splice site variants. Missense variants were prioritized by Mutation Assessor scores (18). Alternative variant prioritization lists were also created using SnpEff/SnpSift^9^. Functional pathway enrichment analysis and annotation were performed with ToppGene (27) and Panther (26).

### Targeted Re-analysis of TCGA Data for RAS Indels

TCGA colorectal tumor whole exome sequencing bam files (n=634) were downloaded from the NIH Genomic Data Commons (GDC) database after appropriate regulatory approval. Indel detection was performed with ScanIndel (14). Briefly, we extracted all reads mapped to *HRAS* (chr11:530,242-537,567), *NRAS* (chr1:114,702,464-114,718,894), and *KRAS* (chr12:25,202,789-25, 252, 931) gene regions (GRCh38). Then BWA-MEM aligned reads were used as input for ScanIndel to find any size of indels with minimum 10x coverage and 1% variant allele frequency thresholds.

### Cell Culture

HEK293T cells (ATCC, CRL-3216) were cultured in complete phenol red free DMEM (Life Technologies, 31053-028, 4.5g/L Glucose, 3.7g/L Sodium Bicarbonate), 2mM L-Glutamine (Life Technologies, 25030081), and 10% FBS (Hyclone, SH30071.03) at 37°C and 5% CO2.

### Transient Transfections

HEK293T cells were seeded at 1.25e5/mL in 12-well plates (Falcon, 353043). After 24 hours, 8μL of Fugene-6 (Promega, E2691) was mixed with various amounts of plasmid DNA and allowed to incubate at room temperature for 15 minutes before addition to cells. After 24 hours, the transfection mixture/media was replaced with complete media and allowed to incubate for another 24 hours.

### NanoBRET Assay

Bioluminescence resonance energy transfer (BRET) is a commonly used imaging assay that uses resonance energy transfer to quantify protein-protein interactions between a bioluminescent donor and fluorescent acceptor(39). For the energy transfer to occur, the donor and acceptor molecules must be within 10 nm of each other and in the correct orientation. The advantage of this system is that full length proteins tagged with the relevant donor acceptor pair can be expressed in cells, and interactions between the proteins of interest can be monitored in live, intact cells. For specific interactions between the donor and acceptor fusion proteins, the BRET ratio increases hyperbolically as a function of increasing acceptor/donor ratio. Saturation is reached when all donor molecules are associated with acceptors. Saturation curves are used to compare the relative affinity of proteins for each other; higher magnitude and steeper increases in signal with lower amounts of acceptor are indicative of greater energy exchange, indicating more molecules are interacting, or that the proximity and geometry of the interaction is favorable for the energy transfer to occur between the donor and acceptor, or both.

In preparation for NanoBRET measurement, transfected cells were incubated in cell dissociation buffer with 0.25% Trypsin-EDTA (Life Technologies, 25200-056). Clumps were then broken up by pipetting with the addition of complete media. After pelleting and discarding the supernatant, cell pellets were dispersed in DMEM with 0.1% FBS, counted, and plated in 384-well white Opti-Plates (Perkin Elmer, 6007290) at a density of 10,000 cells/well. Cells were diluted to 5.0E6 cells/mL in Recovery Cell Culture Freezing Medium (Life Technologies, 12648-010). Halo618 ligand (Promega, N1663) was added and incubated at 37°C for 5 hours. The NanoLuc substrate Furimazine (Promega, N1663) was added at a final concentration of 10μM and incubated for 10 minutes at room temperature. Measurements were made in a Perkin-Elmer Envision plate reader equipped with a 460/80nm band-pass filter (donor) and a 600nm long-pass filter (acceptor).

### NanoBRET Calculations

milliBRET values (mBRET) were calculated by subtracting the A/D ratio, 600nm/460nm, of the donor-only channels from the A/D ratio of the sample channels and then multiplying by 1000.

*mBRET Value* = 1000*((Acceptor channel emission of sample/donor channel emission of sample) – (Acceptor emission channel of donor only/donor channel emission of donor only)) *Z’ value* = 1-3*((SD of Sample + SD of Control)/(mean of Sample – mean of Control)) mBRET values were plotted on GraphPad Prism using the Hyperbola nonlinear equation with concentration of acceptor on the X axis; these curves were used for calculation of BRET_50_ and BRETmax. BRET_50_ is the concentration at which 50% of the maximal mBRET value is achieved, and is considered a relative measure of the affinity of the molecules for each other.

### Western blotting

Western blots were performed using standard experimental approaches to determine induction of MAPK signaling by the ITD mutants as compared to wild-type RAS isoforms. Three replicates were performed for each blot, as described here in detail. After transfection and a 48 hour incubation, HEK293T cells were lysed in a detergent buffer containing 20mM Tris HCl, 150mM NaCl, 1mM EDTA + EGTA, 1% Triton, and Halt protease and phosphatase inhibitor cocktail (Thermo Scientific, no. 78440). Lysates were clarified with centrifugation (16,000xg for 15 minutes at 4°C), protein concentration was analyzed (BCA kit, Thermo Scientific no. 23227), and 25μg protein was loaded per lane onto a Bolt 4-12% 15-well Bis-Tris gel (Invitrogen, no. NW00105BOX). Protein transfer was performed using Thermo Fisher Scientific’s iBlot 2 Dry Blotting System (no. IB21001) and transfer stacks (no. IB23001). After blocking in Odyssey Blocking buffer (LI-COR, no. 927-50003) for one hour at room temperature, membranes were then incubated at 4°C overnight in Odyssey Blocking buffer containing 0.1% Tween 20 and the following antibodies: ERK 1/2 (mouse monoclonal, Cell Signaling Technology no. 4696; 1:1000 dilution), pERK 1/2 (rabbit monoclonal, Cell Signaling Technology no. 4370; 1:2000 dilution), MEK 1/2 (mouse monoclonal, Cell Signaling Technology no. 4694; 1:1000 dilution), pMEK 1/2 (rabbit monoclonal, Cell Signaling Technology no. 9154; 1:1000 dilution), vinculin (mouse monoclonal, Sigma-Aldrich no. V9131; 1:1000 dilution), pan AKT (mouse monoclonal, Cell Signaling Technology no. 2920; 1:1000 dilution), pAKT ser473 (rabbit monoclonal, Cell Signaling Technology no.4060; 1:2000 dilution) and HaloTag (mouse monoclonal, Promega no. G9211; 1:1000 dilution). Membranes were incubated for one hour at room temperature with IRDye secondary antibodies (Goat anti-Mouse 680RD, LI-COR 926-68070; Goat anti-Rabbit 800CW, LI-COR 926-32211; 1:10,000 dilutions). Membranes were washed, and images captures using the LI-COR Odyssey CLx Imaging System. Densitometry analysis was performed with LI-COR Image Studio software. Halotag signal was divided by vinculin signal, and pERK by total ERK, for normalization.

### Cloning, expression, and purification of recombinant proteins

The Gateway Entry clone of *KRAS ITD*, *NF1-GRD* (1198–1530), and *RAF1-RBD* (52–131) were generated by standard cloning methods and incorporated in an upstream tobacco etch virus (TEV) protease cleavage site. Sequence validated entry clones were sub-cloned into pDest-566, a Gateway Destination vector containing a His6 and maltose-binding protein tag (details available at addgene.org/11517) (40). The BL21 STAR (rne131) *E. coli* strain carrying the DE3 lysogen and rare tRNAs (pRare plasmid CmR) was transformed with the expression plasmid (His6-MBP-TEV-protein of interest, AmpR). Proteins were expressed and purified using the procedure described previously (41). Briefly, the expressed proteins of the form His6-MBP-TEV-target were purified from clarified lysates by IMAC, treated with His6-TEV protease to release the target protein, and the target protein separated from other components of the TEV protease reaction by the second round of IMAC. Proteins were further purified by gel-filtration chromatography in buffer containing 20 mM HEPES pH 7.3, 150 mM NaCl, 5 mM MgCl_2_, and 1 mM TCEP. The peak fractions containing pure protein were pooled, flash-frozen in liquid nitrogen, and stored at −80°C.

### Isothermal titration calorimetry measurements

Binding affinities of GMPPNP-bound wild-type and ITD mutant of KRAS (1–169) with NF1-GRD (1098–1530) and RAF1-RBD (52–131) were measured using isothermal titration calorimetry (ITC). Protein samples were prepared by extensive dialysis in a buffer (filtered and degassed) containing 20 mM HEPES (pH 7.3), 150 mM NaCl, 5 mM MgCl_2_, and 1 mM TCEP. For the ITC experiment, 60 μM of KRAS and 600 μM of NF1-GRD or RAF1-RBD were placed in the cell and syringe, respectively. ITC experiments were carried out in a MicroCal PEAQ-ITC instrument (Malvern) at 25 °C using an initial 0.4 μL injection and 18 subsequent injections of 2.2 μL each at 150-second intervals. Data analysis was performed based on a binding model containing “one set of sites” using a nonlinear least squares algorithm incorporated in the MicroCal PEAQ-ITC analysis software (Malvern).

### Crystallization, data collection, and structure determination of GDP-bound RAS ITD

To crystallize *KRAS*-ITD bound to GDP, we carried out crystallization screenings using the sitting-drop vapor diffusion method using sparse matrix screens. The GDP-bound *KRAS*-ITD was crystallized in 200 mM lithium acetate and 2.2 M ammonium sulfate. Crystals were harvested for data collection and cryoprotected with a 30% (v/v) solution of glycerol, before being flash-cooled in liquid nitrogen. The diffraction dataset was collected on 24-ID-E beamline at the Advanced Photon Source (APS), Argonne National Laboratory. Crystallographic datasets were integrated and scaled using XDS (42). Structure of GDP-bound KRAS ITD was solved by molecular replacement using the program Phaser with GDP-bound wild-type KRAS as a search model (43). The initial structure was refined using iterative cycles of manual model building using COOT (44) and refinement using Phenix.refine (45). Placement of ligands was followed by identification of potential sites of solvent molecules by the automatic water-picking algorithm in COOT and Phenix.refine. The positions of these automatically picked waters were checked manually during model building. The data collection and refinement statistics are summarized in Supplemental Table 1. Figures were generated with PyMOL (Schrödinger, LLC).

### Protein Data Bank accession number

The atomic coordinate and structure factor of the GDP-bound KRAS-ITD structure has been deposited in the Protein Data Bank and assigned accession number 6PQ3.

### Study Approval

The patient and family requested that the patient’s body be donated for cancer research upon death. Full written consent was obtained from both parents prior to patient autopsy and harvesting of the patient’s organs and tumors for research. The University of Minnesota Institutional Review Board reviewed the case and provided a research exemption.

### Statistics

Kaplan-Meier estimate of survival in the 2012 TCGA Colorectal Carcinoma cohort(13) was analyzed on the publicly available cBioPortal web interface(46) (47) with the P-value estimated by a log-rank test using default parameters as published. Error bars in all figures represent the data mean +/− SEM.

## Acknowledgements

The authors thank the patient and his parents for donating his remains for the advancement of cancer research, with the hope that this work will be of benefit to future patients. We also thank Dr. Christine Henzler for her guidance regarding bioinformatic analysis, and Michael Franklin for assistance with editorial review of the manuscript. We thank Dom Esposito, Bill Gillette, John Paul Denson, Peter Frank, Zhaojing Meng, Shelley Perkins, Mukul Sherekar, Xiaoying Xe, Nitya Ramakrishnan, Troy Taylor, and Vanessa Wall of the RAS Reference Reagents program (FNLCR) for their help in cloning, protein expression, and purification. We are thankful to the Albert Chan and Timothy Tran for their help with the structural studies. The content of this publication does not necessarily reflect the views or policies of the Department of Health and Human Services, and the mention of trade names, commercial products, or organizations does not imply endorsement by the US Government.

## Research Funding

This research was supported in part by the Department of Laboratory Medicine and Pathology, University of Minnesota (A.C.N.); Eastern Star Scholars, University of Minnesota Masonic Cancer Center (A.C.N.); American Cancer Society Clinical Scientist Development Grant (CSDG-18-139-01-CSM; A.C.N.); the Mezin-Koats Colon Cancer Research Award (E.L.); The Randy Shaver Cancer Research and Community Fund (E.L.); the Litman Family Fund for Cancer Research; a Central Society for Clinical and Translational Research Early Career Development Award (E.L.); Institutional Research Grant #118198-IRG-58-001-52-IRG94 from the American Cancer Society (E.L.); Minnesota Masonic Charities; Minnesota Medical Foundation/University of Minnesota Foundation (E.L.); the Masonic Cancer Center and Department of Medicine, Division of Hematology, Oncology and Transplantation, University of Minnesota (E.L.); and the NIH Clinical and Translational Science KL2 Scholar Award 8UL1TR000114 (to E.L.). Structural work is based upon research conducted at the Northeastern Collaborative Access Team beamlines, which are funded by the National Institute of General Medical Sciences from the National Institutes of Health (P30 GM124165). The Eiger 16M detector on 24-ID-E beam line is funded by a NIH-ORIP HEI grant (S10OD021527). This research used resources of the Advanced Photon Source, a U.S. Department of Energy (DOE) Office of Science User Facility operated for the DOE Office of Science by Argonne National Laboratory under Contract No. DE-AC02-06CH11357. This project was funded in whole or in part with federal funds from National Cancer Institute, NIH Contract HHSN261200800001E.

