## Supplementary material for "*RAS* Internal Tandem Duplication Disrupts GAP-binding to Activate Oncogenic Signaling": Suppl Table 1

### Supplemental Table 1. Data collection and refinement statistics.

|  | GDP-bound KRAS ITD |
| --- | --- |
| Data collection | |
| Wavelength | 0.97918 |
| Resolution range | 41.38 - 1.85 (1.92 - 1.85)* |
| Space group | P 6_3_ |
| Unit cell *a, b, c* (Å)  α, β, γ (°) | 82.77, 82.77, 41.05,  90, 90, 120 |
| Total reflections | 122923 (11078) |
| Unique reflections | 13779 (1329) |
| Multiplicity | 8.9 (8.3) |
| Completeness (%) | 99.75 (98.08) |
| Mean I/sigma(I) | 20.33 (3.18) |
| Wilson B-factor | 26.85 |
| R-merge | 0.07161 (0.6576) |
| CC1/2 | 0.999 (0.93) |
| CC* | 1 (0.982) |
| **Refinement** | |
| Reflections used in refinement | 13779 (1328) |
| Reflections used for R-free | 1379 (132) |
| R-work | 0.1737 (0.2503) |
| R-free | 0.2180 (0.2699) |
| Number of non-hydrogen atoms | 1429 |
| macromolecules | 1345 |
| ligands | 34 |
| solvent | 50 |
| Protein residues | 168 |
| RMS (bonds) | 0.008 |
| RMS (angles) | 1.21 |
| Ramachandran favoured (%) | 96.95 |
| Ramachandran allowed (%) | 3.05 |
| Ramachandran outliers (%) | 0.00 |
| Average B-factor | 41.84 |
| macromolecules | 42.01 |
| ligands | 32.37 |
| solvent | 43.89 |

* Statistics for the highest-resolution shell are shown in parentheses.
